# Genetically Engineered Human Pituitary Corticotroph Tumor Organoids Exhibit Divergent Responses To Glucocorticoid Receptor Modulators

**DOI:** 10.1101/2022.09.07.506977

**Authors:** Saptarshi Mallick, Jayati Chakrabarti, Jennifer Eschbacher, Andreas G. Moraitis, Andrew E. Greenstein, Jared Churko, Kelvin W. Pond, Antonia Livolsi, Curtis Thorne, Andrew S. Little, Kevin C.J. Yuen, Yana Zavros

**Affiliations:** Department of Cellular and Molecular Medicine, University of Arizona College of Medicine, Tucson, Arizona, USA; Department of Neuropathology, Barrow Neurological Institute, Phoenix, Arizona, USA; Corcept Therapeutics, Inc, Menlo Park, California, USA; MilliporeSigma – Life Science, Rockville, Maryland, USA; Department of Neurosurgery, Barrow Neurological Institute, Phoenix, Arizona, USA; Department of Neuroendocrinology, Barrow Neurological Institute, Phoenix, Arizona, USA

**Keywords:** pituitary, organoids, somatostatin, Cushing’s disease, Hedgehog signaling, Mifepristone, Relacorilant

## Abstract

Cushing’s disease (CD) is a serious endocrine disorder attributed to an ACTH-secreting pituitary neuroendocrine tumor (PitNET) that subsequently causes chronic hypercortisolemia. PitNET regression has been reported following treatment with the investigational selective glucocorticoid receptor (GR) modulator relacorilant, but the mechanisms behind that effect remain unknown. Human PitNET organoid models were generated from induced human pluripotent stem cells (iPSCs) or fresh tissue obtained from CD patient PitNETs (hPITOs). Genetically engineered iPSC derived organoids were used to model the development of corticotroph PitNETs expressing USP48 (iPSC^USP48^) or USP8 (iPSC^USP8^) somatic mutations. Organoids were treated with the GR antagonist mifepristone or the GR modulator relacorilant with or without somatostatin receptor (SSTR) agonists pasireotide or octreotide. In iPSC^USP48^ and iPSC^USP8^ cultures, mifepristone induced the predominant expression of SSTR2 with a concomitant increase in ACTH secretion and tumor cell proliferation. Relacorilant predominantly induced SSTR5 expression and tumor cell apoptosis with minimal ACTH induction. Hedgehog signaling mediated the induction of SSTR2 and SSTR5 in response to mifepristone and relacorilant. Relacorilant sensitized PitNET organoid responsiveness to pasireotide. Therefore, our study identified the potential therapeutic use of relacorilant in combination with somatostatin analogs and demonstrated the advantages of relacorilant over mifepristone, supporting its further development for use in the treatment of CD patients.

**HIGHLIGHTS:**

- Cushing disease (CD) is a serious endocrine disorder caused by an adrenocorticotropic hormone (ACTH)-secreting pituitary neuroendocrine tumor (PitNET) that leads to chronic hypercortisolemia
- Mifepristone (Korlym^®^), a non-selective glucocorticoid receptor (GR) antagonist, is an approved treatment for patients with Cushing disease, and competes with the binding of cortisol to the GR as well as the binding of progesterone to the progesterone receptor.
- Relacorilant is an investigational selective GR modulator in development for the treatment of Cushing syndrome that, unlike mifepristone, does not bind to the other hormone receptors.
- Unlike mifepristone, relacorilant does not significantly raise systemic cortisol levels, and cases of PitNET regression with relacorilant have been reported. However, the mechanisms behind these clinical differences remained unknown.
- PitNET organoids were generated from: 1) CRISPR-Cas9 gene editing of patient iPSCs, and 2) CD patient corticotroph PitNETs (hPITOs) and used to compare the diverse effects of mifepristone and relacorilant in a human-relevant model that recapitulates the PitNET microenvironment *in vitro*.
- Mifepristone and relacorilant have different effects on the induction of somatostatin receptor (SSTR) SSTR2 and SSTR5 expression, ACTH secretion and PitNET organoid proliferation and apoptosis.

**BRIEF COMMENTARY:** *Background:* Cushing’s disease (CD), a serious endocrine disorder caused by an adrenocorticotropic hormone (ACTH)-secreting pituitary neuroendocrine tumor (PitNET) leads to chronic hypercortisolemia. Approved for the treatment for CD, Mifepristone (Korlym^®^) is a non-selective glucocorticoid receptor (GR) antagonist with additional competitive binding with progesterone for the progesterone receptor. Relacorilant, an investigational selective GR modulator in development for the treatment of CD, does not bind to the other hormone receptors.

*Translational Significance:* Patient-derived PitNET organoids recapitulate the tumor microenvironment *in vitro.* PitNET organoids revealed the advantages of relacorilant over mifepristone, supporting its further development for use in the treatment of CD.

## 1. Introduction

Cushing’s disease (CD) is a serious endocrine disorder that is caused by an adrenocorticotropic hormone (ACTH)-secreting pituitary neuroendocrine tumor (PitNETs) that leads to excess adrenal cortisol secretion [1–4]. Renamed as PitNETs by the WHO [5], pituitary tumors have been defined as benign, implying that these tumors cause a disease that is not life threatening or harmful to health. However, chronic exposure to excess and unregulated cortisol has wide ranging and detrimental effects on health, including increased stroke rates, diabetes, obesity, depression, anxiety and a threefold increase in the risk of death from cardiovascular disease and cancer [4, 6] [7, 8]. The first-line treatment for CD is pituitary surgery, which is followed by disease remission in only 56% of patients during the 10-year follow-up period post-surgery, even in the hands of an experienced surgeon [9]. Studies have demonstrated that surgical failures and late recurrences of CD are common, and despite multiple treatment regimens, biochemical control still fails in approximately 30% of patients, suggesting that in routine clinical practice, initial and long-term remission is not achieved in a substantial number of patients with CD [7, 10]. These disappointing patient outcomes highlight the need for a better understanding of the pathophysiology of CD to enable the development of novel therapeutic approaches for these patients. Medical therapy is often considered when initial surgery fails to achieve remission, or when the tumor recurs after apparent surgical remission. Approved treatments include pituitary-targeted drugs, adrenal steroidogenic inhibitors, and a non-selective glucocorticoid receptor (GR) antagonist [3, 11, 12]. There is an inverse relationship between disease duration and reversibility of complications associated with the disease, thus emphasizing the importance of identifying an effective medical strategy to rapidly normalize hypercortisolemia/cortisol activity by directly targeting the pituitary adenoma [4, 7, 10]. Standard of care treatments have low efficacy and/or tolerability, making CD a medical therapeutic challenge. For example, mifepristone (Korlym^®^), a non-selective GR antagonist approved by the Food and Drug Administration in 2012 for the treatment of hyperglycemia secondary to endogenous Cushing’s syndrome, including CD, in adults who have failed surgery or are not candidates for surgery, competes with cortisol binding at the GR and significantly improves hyperglycemia secondary to hypercortisolism in patients with CD [13]. However, while mifepristone improves the symptoms of CD, it also raises ACTH (and thus cortisol) levels and exhibits significant side effects in some patients. Through its binding to the progesterone receptor, mifepristone causes side effects including endometrial thickening and vaginal bleeding [13, 14]. Relacorilant is an investigational, selective GR modulator that, unlike mifepristone, has no detectable binding to the progesterone, estrogen and androgen receptors [15, 16]. In a phase 2 study in patients with Cushing’s syndrome, relacorilant reversed the effects of excess cortisol on hypertension and insulin resistance [16], did not significantly raise ACTH or cortisol levels, and has been reported to induce tumor regression in some CD patients with pituitary macroadenomas [17, 18]. Relacorilant is currently under investigation in phase 3 trials for the treatment of patients with endogenous Cushing’s syndrome (GRACE (NCT03697109) and GRADIENT (NCT04308590) studies). The mechanisms driving relacorilant-induced tumor regression, and the role of tumor sensitization to endogenous somatostatin, remain unknown due to the lack of advanced multicellular *in vitro* models that recapitulate the complexity of the PitNET microenvironment. The absence of preclinical models that recapitulate the tumor tissue has prevented us from acquiring the knowledge to develop and implement therapies specifically targeted the tumor with a higher efficacy and tolerability for CD patients.

Pituitary-directed drugs that target the somatostatin receptor (SSTR), expressed on the surface of corticotroph PitNETs, have been used to treat CD [19]. Pasireotide is a somatostatin analogue known to bind to SSTR1, 2, 3, and with highest-affinity binding to SSTR5 [20]. In two large prospective studies, pasireotide monotherapy normalized urinary free cortisol in up to 42% of patients with CD [21, 22]. Combination therapy may show promise whereby pasireotide administered together [23] with cabergoline (a dopamine receptor subtype 2 antagonist) and low-dose ketoconazole (a Hedgehog receptor smoothened (SMO) inhibitor that also inhibits cortisol synthesis) induced biochemical remission in six of eight patients at day 80, increasing the number of patients with a complete response to 88% [21]. However, such treatments are designed without clear understanding and consideration for the receptor dynamics that may change during treatment. Cortisol suppresses somatostatin receptor expression, so GR modulation may affect SSTR abundance and/or sensitivity to endogenous somatostatin. Therefore, while somatostatin receptor subtypes (SSTR) 2 and 5 have been targeted to suppress tumor growth and hormone secretion, the therapeutic benefits of somatostatin analogs are limited in patients with CD.

Given the intriguing preliminary clinical effects of relacorilant on PitNET regression, we sought to identify the mechanism of GR modulation alone or with somatostatin analogs using the PitNET organoids. PitNET organoids were derived from: 1) CRISPR-Cas9 gene editing of patient iPSCs, and 2) CD patient corticotroph PitNETs (hPITOs). These studies demonstrated that SSTR2 and 5 are targets of Hedgehog transcription effector Gli1, and this response is attenuated by activation of the GR pathway. In addition, genetically engineered iPSCs expressing somatic mutations relevant to CD or hPITOs exhibit divergent responses to mifepristone and relacorilant and may explain the clinical observations made in patients.

## 2. Materials and Methods

### 2.1. Generation, Culture and Differentiation of Induced Pluripotent Stem Cells (iPSCs) to Develop PitNET Organoids

Peripheral Blood Mononuclear Cells (PBMCs) from a healthy individual (JCAZ001) were used to generate iPSCs. Six well culture plates coated with 2 mL/well 0.67% Matrigel™ (diluted in E8 media, UA iPSC core, 151169-01) were used to initially culture and expand the iPSCs. The iPSC lines were reprogrammed at the University of Arizona iPSC Core using an established protocol [24]. Once reaching 70% confluency, iPSCs were passaged to Matrigel™ coated 24 well plates at a ratio of 1:8 and grown to 85-90% confluency before beginning the directed differentiation schedule (**Supplemental Figure 1A**) that followed the culture of cells as follows: 1) 0 to 3 with E6 media supplemented with 1% penicillin/streptomycin, 10 μM SB431542 and 5 ng/ml BMP4, 2) at day 3 BMP4 was withdrawn, 3) at day 4 cells were cultured in E6 media, supplemented with 10 μM SB431542, 30 ng/ml human recombinant SHH, 100 ng/ml FGF8b, 10ng/ml FGF18 and 50 ng/ml FGF10, and 4) cells were harvested at day 15 using cold E6 media and resuspended in Matrigel™ (20,000 cells/50 μl Matrigel™). Matrigel™ domes containing iPSCs were plated in culture dishes and overlaid with differentiation media (E6 media supplemented with 10 μM Y-27632, 30 ng/ml human recombinant SHH, 100 ng/ml FGF8b, 10ng/ml FGF18 and 50 ng/ml FGF10, **Supplemental Table 1**), and cultured for a further 15 days at 37°C at a relative humidity of 95% and 5% CO_2_ during which time the 3D organoids developed and used for experiments and analyses. All human iPSC lines were negative for mycoplasma contamination as tested using the Mycoalert Mycoplasma testing kits (LT07-318, Lonza) and no karyotype abnormalities were found (KaryoStat+, Thermo).

### 2.2. CRISPR/Cas9 Gene Editing and Validation of iPSCs

A CRISPR/Cas9 approach was used to genetically engineer iPSCs to express known somatic USP48 and USP8 mutations of CD [25–28]. Mutation sgRNAs for either USP8 (Gene ID: 9101, Double: Y717C (A2150G) & S718P (T2152C)), and USP48 (Gene ID: 84196; M415I (G1245A) and M415V (A1243G), **Supplemental Figure 2A**) were created using the CRISPR Design Tool (SIGMA VC40007) with the assistance of the SIGMA Bioinformatics Support Team. Single guide RNAs (sgRNAs) targeted specific restriction sites whereby USP8 destroyed a BseRI site and created a BstBI site, and USP48 created a SpeI site. SgRNAs were cloned into a LV01/pLV-U6g-EPCG expression vector (expressing sgRNA and Cas9), and combined with Cas9 protein (SIGMA E120030), warm Opti-MEM^TM^ I Reduced Serum Medium (Thermo Fisher Scientific 31985-070) and *Trans*IT-CRISPR (SIGMA T1706). These *Trans*IT-CRISPR Cas9 Ribonucleotide complexes were then added dropwise to their respective iPSC plates and incubated for 48 hours before starting the initiation of the differentiation schedule.

Successful gene editing of the iPSCs was validated using restriction fragment length polymorphism (RFLP) analysis. DNA was extracted from iPSCs using the QIAamp DNA Micro Kit (Qiagen 56304) according to manufacturer’s protocol, and the concentration was measured using a NanoDrop™ One/OneC Microvolume UV-Vis Spectrophotometer (Thermo Fisher Scientific ND-ONE-W). Amplification of 300 ng of DNA was performed using KOD Hot Start Master Mix (Sigma-Aldrich 71842), forward and reverse primers of the mutations (USP8 Forward: TTG AAG TTT ATC GCC ATT TTA TTC G, Reverse: GCT GGT ATA GCC ATC CAC AGA A; USP48 Forward: TGT TGT TCT AGG TTT GTT CCC CA, Reverse: TCA CCA ACA TTC ACC TTA TGG AA). The samples were amplified in a thermal cycler (Applied Biosystems Veriti™ 96-Well Thermal Cycler, Thermo Fisher Scientific 4375786) with the following temperature cycles: 95° for 2 minutes, 40 cycles of 95°C for 20 seconds, 48°C for 10 seconds, 70°C for 1 minute. 6X Purple Gel Loading Dye (NEB B7025S) was added to the amplicon and the PCR products were loaded onto a 1% agarose gel. After electrophoresis, the bands were visualized using a UV imager, extracted using the GenElute™ Gel Extraction Kit (SIGMA NA1111) and purified with the GenElute^™^ PCR Clean-Up Kit (SIGMA NA1020). 300 ng of the eluted DNA was digested overnight at 37°C with the restriction enzymes including USP8 – BstBI and USP48 – SpeI. Restriction fragments were visualized on a 2.5% agarose gel alongside undigested DNA (**Supplemental Figure 2B, C**).

### 2.3. Generation and Culture of Human PitNET Tissue Organoids (hPITOs)

Patients undergoing planned transsphenoidal surgery for PitNETs were identified in the outpatient neuroendocrinology and neurosurgery clinics and collected by the St. Joseph’s Hospital and Barrow Neurological Institute Biobank under the collection protocol PHXA-05TS038, and outcomes data protocol PHXA-0004-72-29 with approval of the Institutional Review Board (IRB) and patient consent. De-identified samples were shipped in collection and transport media (**Supplemental Table 2**) to the Zavros laboratory (University of Arizona) for processing within 24 hours of collection. Resected PitNET tissues were washed with DPBS supplemented with 1X Penicillin/Streptomycin, 1X Kanamycin and 1X Amphotericin/Gentamycin, minced into smaller pieces, and incubated with digestion buffer (DMEM/F12 supplemented with 0.4% Collagenase 2, 0.1% Hyaluronic Acid, 0.03% Trypsin). The tissue was incubated with Accutase (Thermo Fisher Scientific) for a further 5 minutes, and the cells were pelleted and washed with DPBS supplemented with antibiotics. Dissociated cells were then seeded as Matrigel™ domes in culture dishes and overlaid with growth media (**Supplemental Table 3**). Media was replaced every 3 days and organoids were harvested for downstream experiments or passaged after 15 days of culture (**Supplemental Figure 1B**).

### 2.4. Immunofluorescence

iPSCs were stained for pituitary specific hormones including luteinizing hormone (LH), follicle-stimulating hormone (FSH), growth hormone (GH), prolactin (PRL), ACTH, cytokeratin 8/18 (CAM5.2), pituitary transcription factor PIT-1 and neuroendocrine marker Synaptophysin (**Supplemental Table 4**). Cultures were immunostained for proliferation marker EdU (Click-IT EdU Alexa Fluor 555 Imaging Kit, Thermo Fisher Scientific C10338). Antibody dilutions were determined based on manufacturer guidelines and antibody titration (**Supplemental Table 4**). Staining was conducted according to previously published protocols [29, 30]. Briefly, organoids and iPSCs grown in 8-well glass bottom chamber slides were fixed using 3.7% formaldehyde, washed with DPBS, permeabilized with 0.5% Triton X-100 (SIGMA X-100, PBS-T, 20 min at room temperature), washed with 0.01% PBS-T, and blocked with 2% donkey serum (Jackson Immuno Research 017-000-121) for 1 hour at room temperature. After blocking, primary antibodies were added and incubated overnight at 4°C. Slides were washed with 0.01% PBS-T and incubated with secondary antibodies (**Supplemental Table 4**) with Hoechst (Thermo Fisher Scientific H1399) for 1 hour at room temperature. Organoids were washed and stored in PBS at 4°C until microscopic analysis. High content confocal microscopy was performed using the Nikon Ti2-E Inverted Microscope (with a Crest X-Light V2 L-FOV Spinning Disk Confocal) at the University of Arizona Cancer Center, or a ZEISS LSM880 34 channel Laser Scanning Confocal Microscope at University of Arizona Marley Imaging Core. Fluorescence intensity was quantified as the percentage of marker-positive cells versus total cells, using Nikon Elements Software (Version 5.21.05). Proliferation for each iPSC line was quantified as the percentage of EdU-positive cells versus total cells.

### 2.5. Nuclear Morphometric Analysis (NMA)

Nuclear Morphometric Analysis (NMA) using treated organoids was performed based on a published protocol that measures cell viability based on the changes in nuclear morphology of the cells using nuclear stain Hoechst or DAPI [31]. Images of organoid nuclei were analyzed using the ImageJ Nuclear Irregularity Index (NII) plugin for key parameters that included cell area, radius ratio, area box, aspect, and roundness. Using the published spreadsheet template [31], the NII of each cell was calculated with the following formula: NII = Aspect – Area Box + Radius Ratio + Roundness. The area vs NII of vehicle-treated cells were plotted as a scatter plot using the template and considered as the normal cell nuclei. The same plots were generated for each condition, and the NII and area of treated cells were compared to the normal nuclei, and classified as one of the following NMA populations: Normal (N; similar area and NII), Mitotic (S; similar area, slightly higher NII), Irregular (I; similar area, high NII), Small Regular (SR; apoptotic, low area and NII), Senescent (LR; high area, low NII), Small Irregular (SI; low area, high NII), or Large Irregular (LI; high area, high NII). Cells classified as SR exhibited early stages of apoptosis, and cells classified as either I, SI or LI exhibited significant nuclear damage. The percentage of cells in each NII classification category were calculated and plotted as a histogram using GraphPad Prism.

### 2.6. Immunohistochemistry (IHC)

PitNET organoids were fixed in 4% paraformaldehyde and embedded in Epredia™ HistoGel™ Specimen Processing Gel according to the Manufacturers protocol (Fisher Scientific HG-4000-012). HistoGel embedded organoids were then paraffin embedded and sectioned on slides at a 5-micron thickness. After deparaffinization and antigen retrieval (Antigen Unmasking Solution, Vector Laboratories H-3300-250), endogenous peroxidase activity was blocked using 0.3% hydrogen peroxide/methanol for 20 min. Slides were blocked with 20% normal goat or horse serum for 20 min at room temperature and incubated with primary antibodies overnight at 4 °C (**Supplemental Table 4**). Slides were incubated with biotinylated anti-mouse or anti-rabbit secondary antibodies for 30 minutes. Color was developed with 3,3′-diaminobenzidine (DAB) using the DAB substrate VECTASTAIN Elite Mouse (Vector Laboratories PK-6102) or Rabbit (Vector Laboratories PK-6101). Immunohistochemical slides were dehydrated and mounted using Permount (Fisher Scientific), and images were viewed and captured under light microscopy (Olympus BX60 with Diagnostic Instruments “Spot” Camera; Tokyo, Japan).

### 2.7. Organoid Drug Treatments and Dose Responses

Sonic Hedgehog (SHH) was withdrawn from the organoid growth media 16 hours prior to treatment. The experimental treatment groups included: vehicle (Veh), mifepristone (Mife, 500nM, Selleckchem S2606), GANT61 (5µM, Stemcell Technologies 73692), ketoconazole (Keto, 10µM, Selleckchem S1353), pasireotide (Pas, 100nM, Target Mol TP2207-SB001), octreotide (Oct, 100nM, Selleckchem P1017), Mife plus GANT61, Mife plus Keto, Mife plus Pas, Relacorilant (Rela, 500nM, Corcept Therapeutics CORT125134), Rela plus Pas, Rela plus Oct, or dexamethasone (Dexa, 100nM, SIGMA S4902). In the combination treatments, cultures were pretreated with the GANT61, Keto, Pas or Oct for 2 hours prior to treatment with Mife or Rela. PitNET tissue derived organoids hPITO36, hPITO37, hPITO38, hPITO39, and hPITO40 were treated with the following experimental conditions: Vehicle (1), pasireotide (2), relacorilant (3), relacorilant & pasireotide (4), mifepristone (5), mifepristone & pasireotide (6). The IC50 values used for each hPITO line for each drug were determined based on dose responses (**Supplemental Figure 3)**. All organoid cultures were treated for 48 hours prior to analysis.

### 2.8. ACTH Enzyme-Linked Immunosorbent Assay (ELISA)

Organoid conditioned media was collected and ACTH secretion using an ELISA according to the manufacturer’s protocol (Novus Biologicals Human ACTH ELISA Kit NBP2-66401). The optical density was measured at a wavelength of 450nm, and ACTH concentration (pg/mL) was interpolated using a standard curve with a 4-parameter logistic regression analysis using GraphPad Prism (v9.3.1).

### 2.9. Quantitative RT-PCR (qRT-PCR)

RNA was extracted from PitNET tissue or iPSC generated organoids using TRI Reagent® (Molecular Research Center Inc TR118) and the High Capacity cDNA Reverse Transcription Kit (Applied Biosystems, Foster City, CA) was used for cDNA synthesis of RNA following the recommended protocol. For each sample, 60 ng RNA was reverse transcribed to yield approximately 2 μg total cDNA that was then used for the qRT-PCR. Pre-designed real-time polymerase chain reaction assays were purchased for the following genes: TPit (Thermo Fisher Scientific 4331182 Hs00193027), SF1 (Thermo Fisher Scientific 4331182 Hs00610436), SSTR2 (Thermo Fisher Scientific 4331182 Hs00265624_s1), SSTR5 (Thermo Fisher Scientific 4331182 Hs00990407_s1), POMC (Thermo Fisher Scientific 4331182 Hs01596743_m1), hPRT (Thermo Fisher Scientific 4331182 Hs02800695_m1), and FKBP5 (Thermo Fisher Scientific 4331182 Hs01561006_m1). Polymerase chain reaction amplifications were performed in a total volume of 20 μL containing 20X TaqMan Expression Assay primers, 2X TaqMan Universal Master Mix (Applied Biosystems, TaqMan® Gene Expression Systems), and cDNA template. Each polymerase chain reaction amplification was performed in triplicate wells in a StepOne Real-Time PCR System (Applied Biosystems) by using the following conditions: 50°C 2 minutes, 95°C 10 minutes, 95°C 15 seconds (denature) and 60°C 1 minute (anneal/extend) for 40 cycles. Fold change was calculated as the following: (C_t_–C_t high_) = n _target_, 2^ntarget^/2^nHPRT^ = fold change where C_t_ = threshold cycle. The results were expressed as average fold change in or differential gene expression relative to control, with HPRT used as an internal control according to Livak and Schmittgen [32].

### 2.10. Spectral Flow Cytometry Using the Cytek® Aurora

The multicolor flow cytometry antibody panel was designed using the Cytek® Full Spectrum Viewer online tool to calculate the similarity index. Organoids were harvested in cold Serum-Free Defined Medium media and incubated with Zombie (BioLegend 423107, 1:100) for 15 minutes in the dark at room temperature, followed by incubation with fluorochrome-conjugated/unconjugated primary surface or cytoplasmic antibodies (**Supplemental Table 5**) at 4°C for 30 minutes. Cells were then washed with Cell Staining Buffer (BioLegend 420-201) and incubated at 4°C for 30 minutes with secondary antibodies (**Supplemental Table 5**). Cells were fixed using Cytofix/Cytoperm^TM^ Fixation/Permeabilization Solution (BD Biosciences 554714) and washed with Fixation/Permeabilization wash buffer according to the recommended protocol. Intracellular markers were then labeled with fluorochrome-conjugated/unconjugated primary antibodies at 4°C for 30 minutes, washed and incubated with secondary antibodies at 4°C for 30 minutes (**Supplemental Table 5**). UltraComp eBeads^TM^, Compensation Beads (Thermo Fisher Scientific 01-2222-42) were stained with the individual antibodies and used as single stain control for compensation and gating. For Zombie dye single cell control, ArC™ Reactive Beads (Thermo Fisher Scientific A10346A) were stained with Zombie and ArC™ negative beads (Thermo Fisher Scientific A10346B). Data was acquired using the Cytek® Aurora, analyzed using Cytobank Premium software (Beckman Coulter).

### 2.11. Statistical Analysis

Data collected from each individual study used 4 *in vitro* experimental or biological replicates unless otherwise stated. Results were analyzed as the mean ± the standard error of the mean (SEM), and data represented as violin or scatter plots, and significance tested using Microsoft Excel and GraphPad Prism (v9.3.1).

## 3. Results

### 3.1. IPSC-derived pituitary organoids expressing somatic mutations reveal corticotroph PitNET pathology consistent with CD

Extensive research has revealed the role of somatic mutations in the development of CD adenomas [27, 33]. To study these mechanisms, we generated human pituitary organoids developed from iPSCs. With reference to a published protocol using embryonic stem cells [34] together with the knowledge of the growth factor and genetic regulation of pituitary gland development [35], we optimized our approach for the development of pituitary organoids from human blood-derived iPSCs for gene editing (**Supplemental Figure 1A**). The iPSC lines were generated to express known somatic mutations in iPSC*^USP48^* (Gene ID: 84196; M415I (G1245A) and M415V (A1243G)), or iPSC*^USP8^* (**Supplemental Figure 1A, Supplemental Figure 2A**) [25, 33, 36–41]. USP48 mutations included either M415V or M415I, designating iPSC^USP48MV^ and iPSC^USP48MI^ respectively. Successful gene editing was validated using primer specific visualization of a change in the restriction pattern at the site of interest (**Supplemental Figures 2B, C**). Brightfield images showed morphological variations between the control (iPSC^ctrl^) and mutant iPSCs as early as day 4 and at day 15 (**Supplemental Figure 1A**). Morphological changes and proliferation were obvious when iPSCs were embedded in Matrigel™ to generate organoids (**Supplemental Figure 1A**).

Expression of PIT1 (pituitary-specific positive transcription factor 1), ACTH (adrenocorticotropic hormone), GH (growth hormone), FSH (follicle-stimulating hormone), LH (luteinizing hormone), PRL (prolactin) and synaptophysin (synapto) with co-stain Hoechst (nuclei, blue) was measured by immunofluorescence using chamber slides collected at days 4 (D4, **Supplemental Figure 4**) and 15 (D15, **Figure 1**) of the differentiation schedules. While pituitary tissue that was differentiated from control iPSCs (iPSC^ctrl^) expressed all major hormone-producing cell lineages (**Figure 1A**), there was a significant increase in the expression of ACTH and synaptophysin with a concomitant loss of PIT1, GH, FSH, LH and PRL in iPCSs expressing mutated USP48 and USP8 (**Figures 1B, C, D**). Immunofluorescence of iPSCs collected at day 4 of the differentiation schedule revealed no expression of PIT1, ACTH, GH, FSH, LH and PRL in iPSC^ctrl^ (**Supplemental Figure 4A**). Unexpectedly, ACTH expression was increased in iPSC^USP48^ at day 4 (**Supplemental Figures 4B, C**).

**Figure 1:**
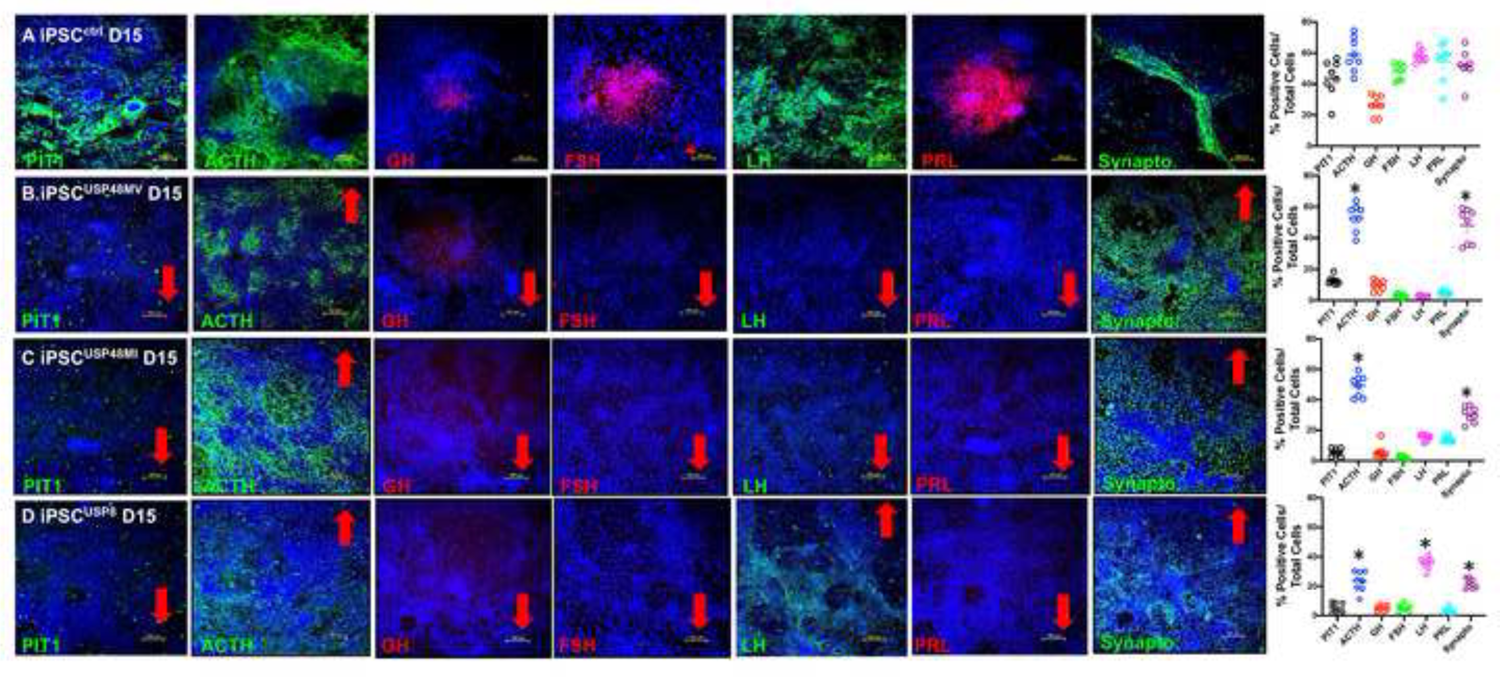
Expression pattern of pituitary hormone-producing cell lineages in iPSCs differentiated to pituitary organoids. Expression of PIT-1 (green), ACTH (green), GH (red), FSH (red), LH (green), PRL (red) and synaptophysin (synapto, green) with co-stain Hoechst (nuclei, blue) was measured by immunofluorescence using chamber slides collected at day 15 (D15) of the differentiation schedule of **(A)** control iPSCs (iPSC^ctrl^) and iPSCs expressing **(B)** USP48M415V (iPSC^USP48MV^), **(C)** USP48M415I (iPSC^USP48MI^) and **(D)** USP8 (iPSC^USP8^) mutations. Red arrows highlight the increased expression of ACTH and synaptophysin with the concomitant loss of PIT1, GH, FSH, LH and PRL in iPCSs expressing somatic mutations USP48 and USP8. Quantification of the percentage of positive cells is shown in the dot plots for each iPSC line.

Immunohistochemistry of formalin fixed and paraffin embedded iPSC^ctrl^, iPSC^USP48^, and iPSC^USP8^ expressed CAM5.2, T-PIT, synaptophysin and ACTH (**Figure 2A**). The expression pattern of corticotroph PitNET specific markers was analyzed using Cytek™ Aurora spectral flow cytometry (**Figure 2B**). The location of cells that were found in each cluster based on the highly expressed antigens are shown in the representative heatmap t-SNE maps (**Figure 2B**). Compared to iPSC^ctrl^ organoids, iPSC^USP48^ and iPSC^USP8^ contained higher numbers of cells expressing T-Pit cell lineage and ACTH/SSTR2/SSTR5 expressing cells (representative t-SNE map of iPSC^USP48MV^ shown in **Figure 2B**). In addition, iPSC^ctrl^ organoids expressed a higher percentage of Pit1 cell lineage, GH and PRL positive cells (**Figure 2B**).

The differential expression of the major transcription factors was correlated with the classification of PitNETs based on immunofluorescence (**Figure 2C, D**). The iPSC^ctrl^ line expressed transcription factors TPit (*TBX19*), PIT1 (*POU1F1*) and *SF1* that corresponded to the differentiation of corticotrophs, somatotrophs and gonadotrophs as documented by positive immunofluorescence staining (**Figures 2C, D**). The iPSC lines expressing mutations in USP48MV exhibited significantly elevated transcription factor T-Pit consistent with the skewed differentiation in the corticotroph cell lineage (**Figures 2C, D**). Transcription factors PIT1 (somatotrophs) and SF1 (gonadotroph) were significantly decreased in the cultures of iPSC lines expressing USP48MV compared to iPSC^ctrl^ (**Figure 2C, D**).

**Figure 2:**
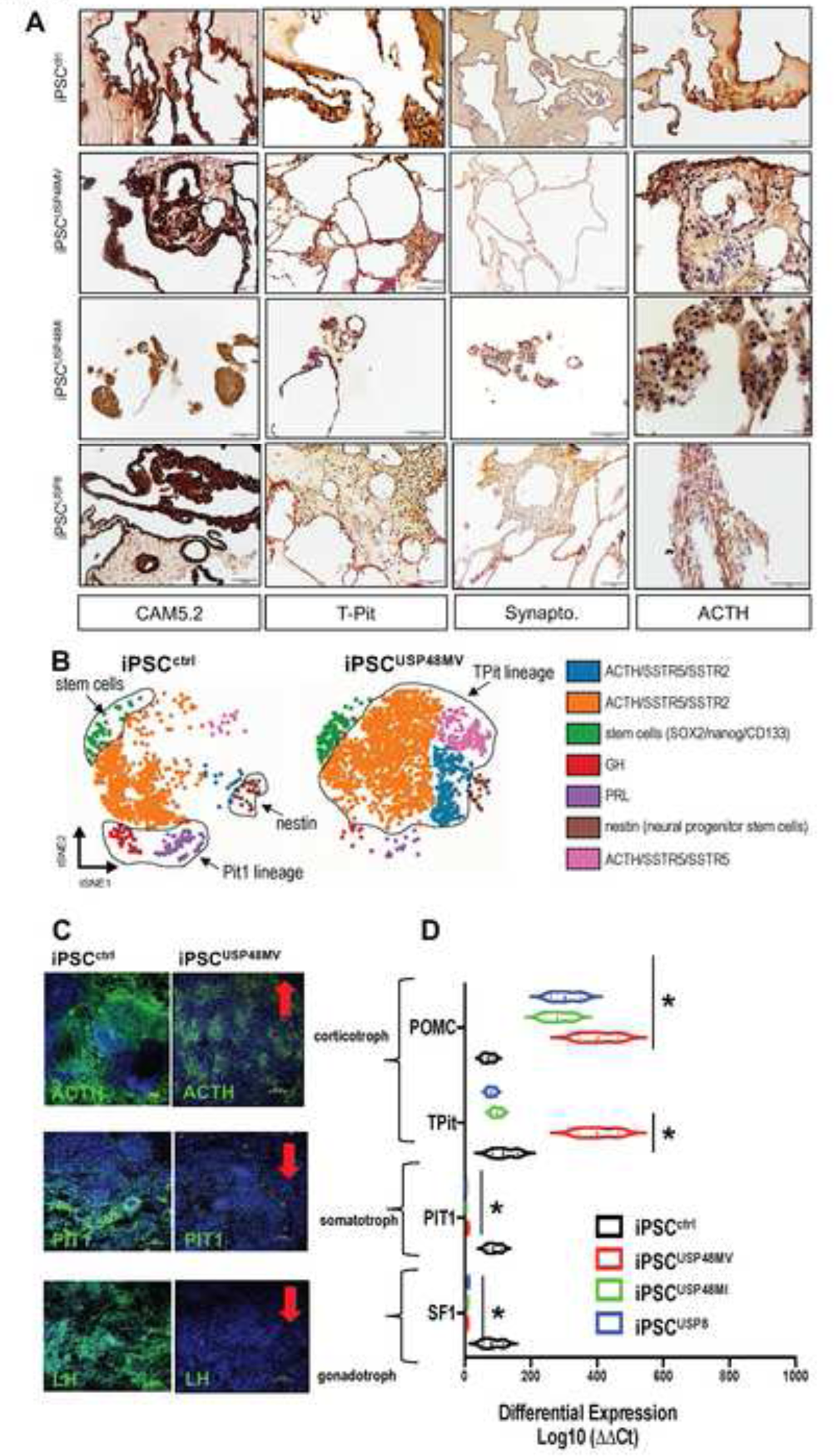
Expression of cell linages and transcription factors in iPSC generated pituitary tumor organoids. **(A)** Immunohistochemistry using FFPE sections prepared from iPSC^ctrl^, iPSC^USP48MV^, iPSC^USP48MI^, and iPSC^USP8^ organoids stained with antibodies specific for CAM5.2, T-Pit, Synaptophysin and ACTH. **(B)** viSNE maps showing concatenated flow cytometry standard files for iPSC^ctrl^, iPSC^USP48MV^, iPSC^USP48MI^, and iPSC^USP8^ organoids 30 days post-directed differentiation. Maps define spatially distinct cell populations using pituitary specific cell lineage, stem cell and transcription factor markers between iPSC^ctrl^ organoids and mutant lines. **(C)** Immunofluorescence staining of ACTH, PIT1 and LH expression in iPSC^ctrl^ and iPSC^USP48MV^ cultures 15 days post-directed differentiation. **(D)** Differential expression of transcription factors TPit, PIT1 and SF1 and POMC were measured by qRT-PCR. *p<0.05 compared to iPSC^ctrl^ organoids, n = 3 individual experimental replicates.

Culture conditioned media was collected during the differentiation schedule of the iPSCs and analyzed for ACTH secretion by ELISA (**Figure 3A**). Compared to control lines, iPSC lines expressing mutated *USP8* and *USP48* secreted significantly greater concentrations of ACTH earlier in the differentiation schedule (**Figure 3A**). Differentiated iPSCs were then collected at D15 of the culture schedule and embedded into Matrigel™. Proliferation was quantified by EdU uptake of the cells within the pituitary organoids (**Figure 3B**). Compared to control pituitary organoids iPSC^ctrl^, organoids expressing mutations in USP8 (iPSC^USP8^), and USP48 (iPSC^USP48MV^ and iPSC^USP48MI^) expressed significantly elevated Edu+ proliferating cells (**Figure 3C**). Collectively, these data show that iPSCs genetically engineered to express somatic mutations relevant to the development of CD exhibit corticotroph PitNET pathology and function *in vitro*.

**Figure 3:**
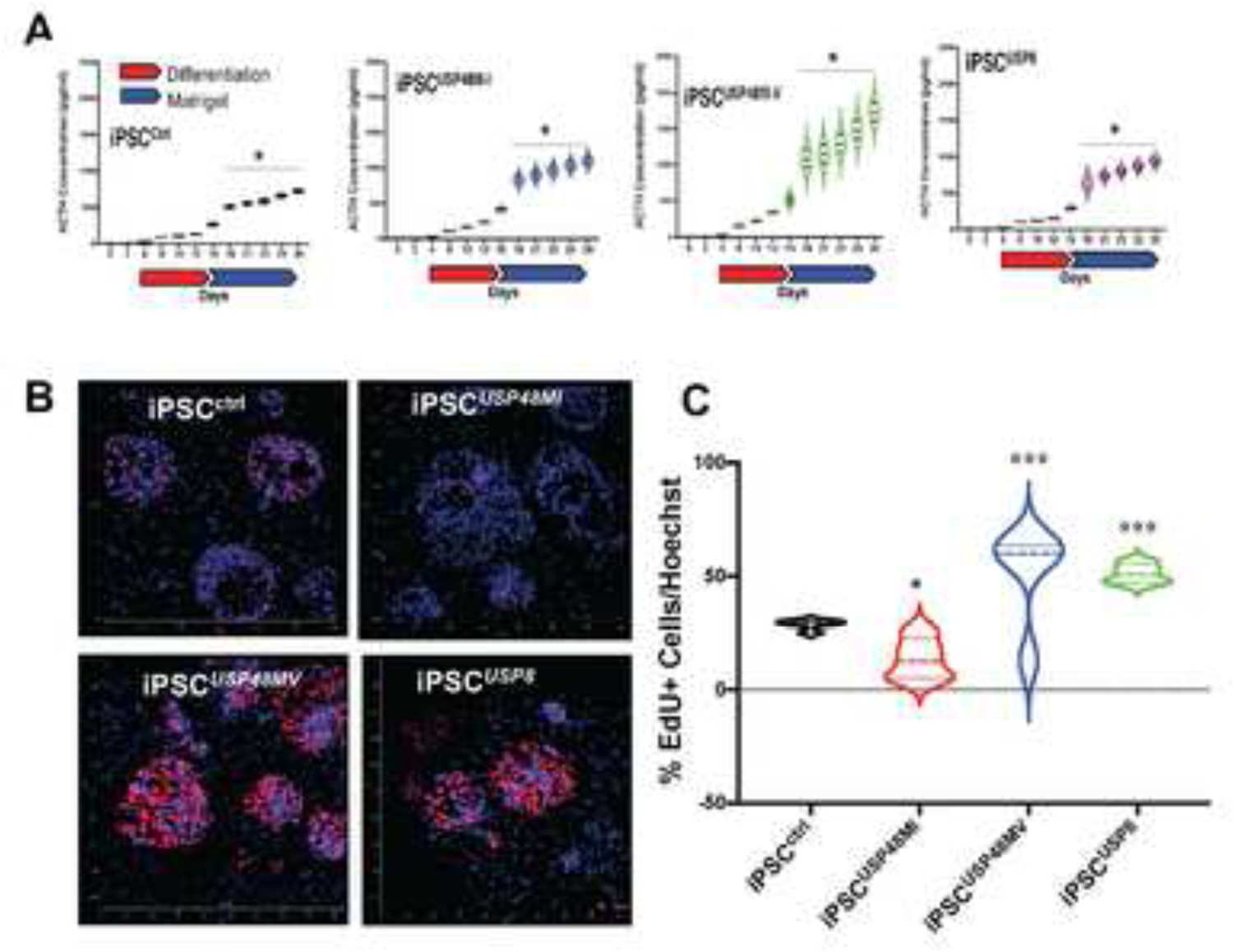
Proliferation and ACTH Secretion During Directed Differentiation of iPSC organoids. **(A)** An ELISA was performed using conditioned media collected during the differentiation schedule from iPSC^ctrl^ and iPSC mutant cultures for the measurement of ACTH secretion (pg/ml). **(B)** Immunofluorescence images of expression of EdU in iPSC^ctrl^, iPSC^USP48MI^, iPSC^USP48MV^, and iPSC^USP8^ organoids. **(C)** Quantification of EdU positive cells of iPSC^ctrl^ and mutant organoids. * p< 0.05, *** p<0.0001 compared to iPSC^ctrl^ organoids, n = 4 individual organoids quantified per culture.

### 3.2. Genetically engineered iPSCs expressing somatic mutations relevant to CD identify differential effects of relacorilant and mifepristone on SSTR expression, and tumor cell proliferation and apoptosis

The treatment of iPSC organoids with mifepristone or relacorilant resulted in a significant induction in the expression of SSTR2 and 5 (**Figure 4A, B**). Mifepristone led to a significantly greater induction in SSTR2 expression when compared to relacorilant (**Figure 4A**). In contrast, relacorilant induced greater SSTR5 expression (**Figure 4B**).

**Figure 4:**
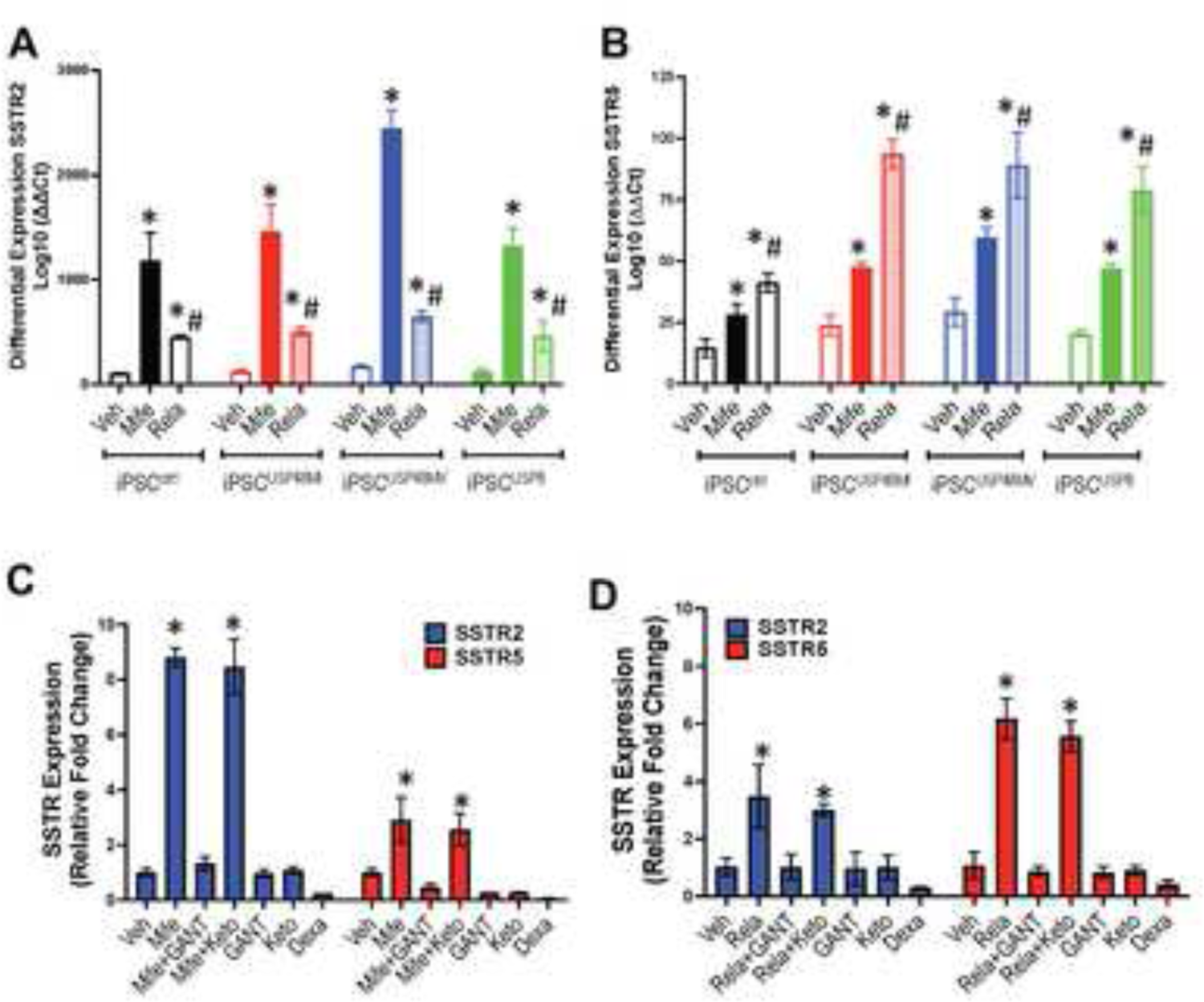
Differential expression of SSTR2 and SSTR5 in iPSC generated pituitary tumor organoids in response to Mifepristone and Relacorilant. Differential expression of (A) SSTR2 and (B) SSTR5 in iPSC^ctrl^, iPSC^USP48MI^, iPSC^USP48MV^ and iPSC^USP8^ organoids in response to vehicle (Veh), mifepristone (Mife) or relacorilant (Rela). *p<0.05 compared to vehicle-treated organoids, #p<0.05 compared to mifepristone treated organoids, n = 3 experimental replicates/organoid line. (C) Differential expression of SSTR2 and SSTR5 in iPSC^ctrl^ organoids treated with vehicle (Veh), mifepristone (Mife), GANT61 (GANT), Mife+GANT, ketoconazole (Keto), Mife+Keto, or dexamethasone (Dexa). *p<0.05 compared to iPSC^ctrl^ organoids treated with Veh, n = 3 experimental replicates/organoid line. (D) Differential expression of SSTR2 and SSTR5 in iPSC^ctrl^ organoids treated with vehicle (Veh), relacorilant (Rela), GANT61 (GANT), Rela+GANT, ketoconazole (Keto), Rela+Keto, or dexamethasone (Dexa). *p<0.05 compared to iPSC^ctrl^ organoids treated with Veh, n = 3 experimental replicates/organoid line.

To identify the role Hedgehog signaling as a mediator of mifepristone or relacorilant regulated SSTR expression, iPSC^ctrl^ organoids were used in combination with inhibitors GANT61 (GLI inhibitor) and ketoconazole (a SMO inhibitor that also inhibits cortisol synthesis) (**Figure 4C, D**). Mifepristone significantly induced the differential expression of both SSTRs 2 and 5, and this increase was reduced with GANT61 pretreatment of the organoid cultures (**Figure 4C**). In contrast to the inhibitory effect of GANT61 on mifepristone-induced SSTR expression, ketoconazole pretreatment had no effect on this induction (**Figure 4C**). GANT61 and ketoconazole alone had no effect on the differential expression of SSTRs 2 and 5, while dexamethasone, a known GR agonist, significantly decreased expression of these receptors (**Figure 4C**). Similar responses were observed with relacorilant with or without GANT61 and ketoconazole (**Figure 4D**). These results suggest that mifepristone and relacorilant induce SSTR2 and SSTR5 via a SMO-independent non-canonical Hh signaling pathway.

Analysis of proliferation using EdU uptake in iPSC^ctrl^ and iPSC^USP48MV^ organoid cultures showed that Mife induced a significant increase in cell proliferation (**Figure 5A, B** and **Figure 6A, B**). While the proliferative response to Mife was inhibited by pretreatment with Oct, Pas had no effect (**Figure 5A, B** and **Figure 6A, B**). While a similar response was observed in Rela treated cultures with pretreatment with Pas or Oct, Rela alone did not induce organoid proliferation (**Figure 5A, B** and **Figure 6A, B**). The nuclear irregularity index (NII) was measured based on the quantification of the morphometric changes in the nuclei in response to the treatment groups in the iPSC^ctrl^ (**Figure 5C-F, Supplemental Figure 5**) and iPSC^USP48MV^ organoids (**Figure 6C-F, Supplemental Figure 6**). While Rela induced a significant expression pattern of nuclear morphology consistent with increased apoptotic cells in the iPSC^USP48MV^ cultures (**Figure 6C-F, Supplemental Figure 6**), this was not observed in the iPSC^ctrl^ organoids (**Figure 5C-F, Supplemental Figure 5**). The magnitude of the apoptotic response to Rela was not observed with Mife in either the iPSC^ctrl^ (**Figure 5C-F, Supplemental Figure 5**) or iPSC^USP48MV^ (**Figure 6C-F, Supplemental Figure 6**) organoids.

**Figure 5:**
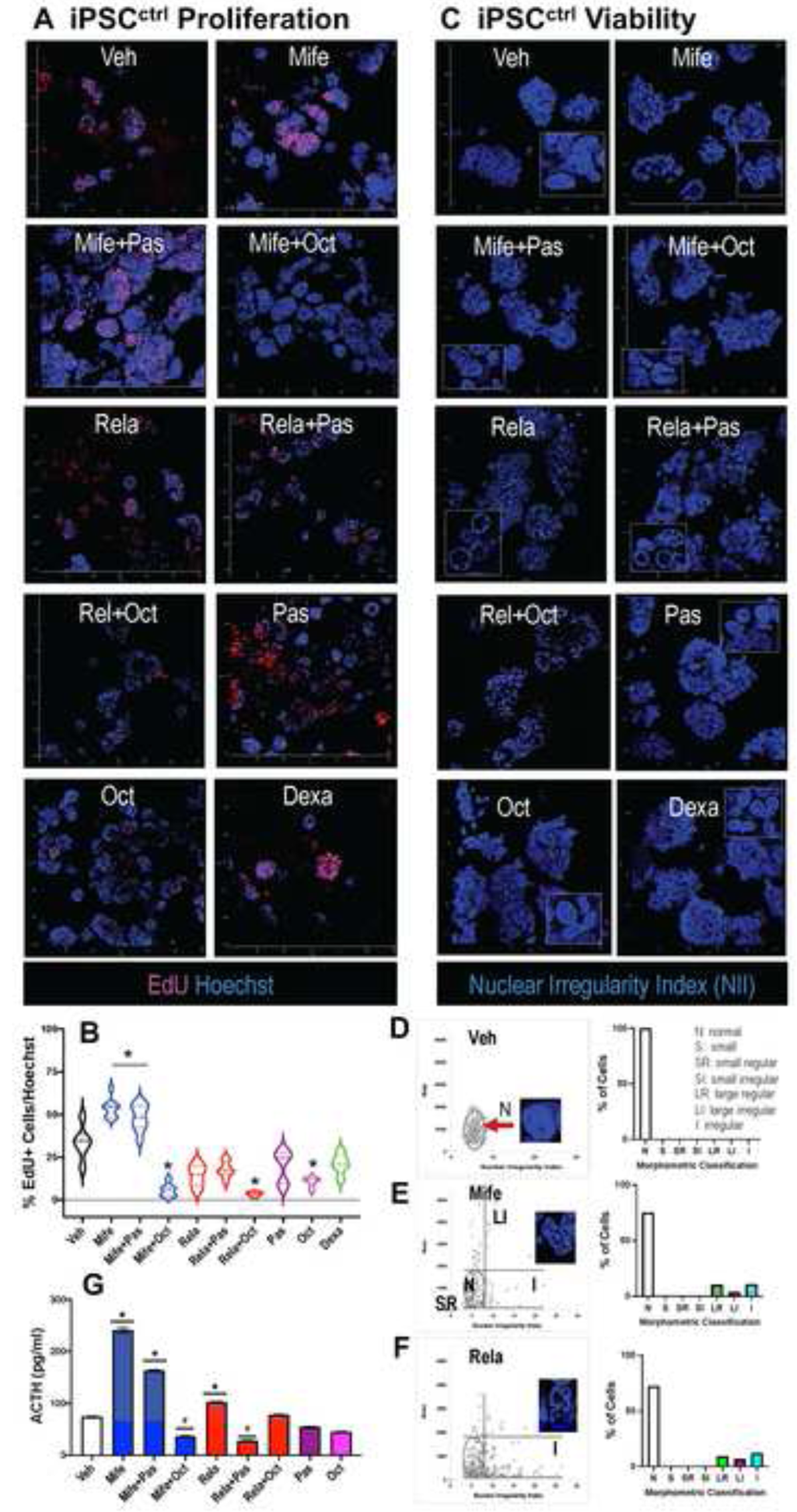
Changes in pituitary tumor cell proliferation, viability and ACTH secretion in iPSC^ctrl^ organoids in response to mifepristone and relacorilant. (A) Immunofluorescence images of EdU expression in iPSC^ctrl^ organoids in response to vehicle (Veh), mifepristone (Mife), pasireotide (Pas), octreotide (Oct), relacorilant (Rela), Mife+Pas, Mife+Oct, Rela+Pas, Rela+Oct, dexamethasone (Dexa). (B) Quantification of EdU positive cells of iPSC^ctrl^ and mutant organoids. *p< 0.05 compared to iPSC^ctrl^ organoids, n = 4 individual organoids quantified per culture. (C) Representative Hoescht staining of iPSC^ctrl^ organoids in response to experimental treatments for the calculation of nuclear irregularity index (NII). Nuclear morphometric analysis of iPSC^ctrl^ organoids in response to experimental treatments with quantification shown for (D) Veh, (E) Mife, or (F) Rela treatments. Morphometric classification of NII was based on the normal (N), small (S), small regular (SR), short irregular (SI), large regular (LR), large irregular (LI) and irregular (I) nuclear morphology. (G) An ELISA was performed using conditioned media collected from iPSC^ctrl^ cultures in response to treatments for the measurement of ACTH secretion (pg/ml). *p<0.05 compared to Veh treatment, #p<0.05 compared to Mife or Rela alone, n = 4 individual organoids quantified per culture.

Within the iPSC^ctrl^ and iPSC^USP48MVl^ organoid cultures, the magnitude of ACTH secretion induced by mifepristone was significantly greater than the effect of relacorilant (**Figure 5G and 6G**). Pas and Oct significantly reduced ACTH secretion in response to mifepristone, although the inhibition by octreotide was greater that pasireotide (**Figure 5G and Figure 6G**). Pasireotide reduced hormone secretion in combination with relacorilant at a greater magnitude compared to that of octreotide plus relacorilant in iPSC^ctrl^ and iPSC^USP48MV^ organoid cultures (**Figure 5G and Figure 6G**).

**Figure 6:**
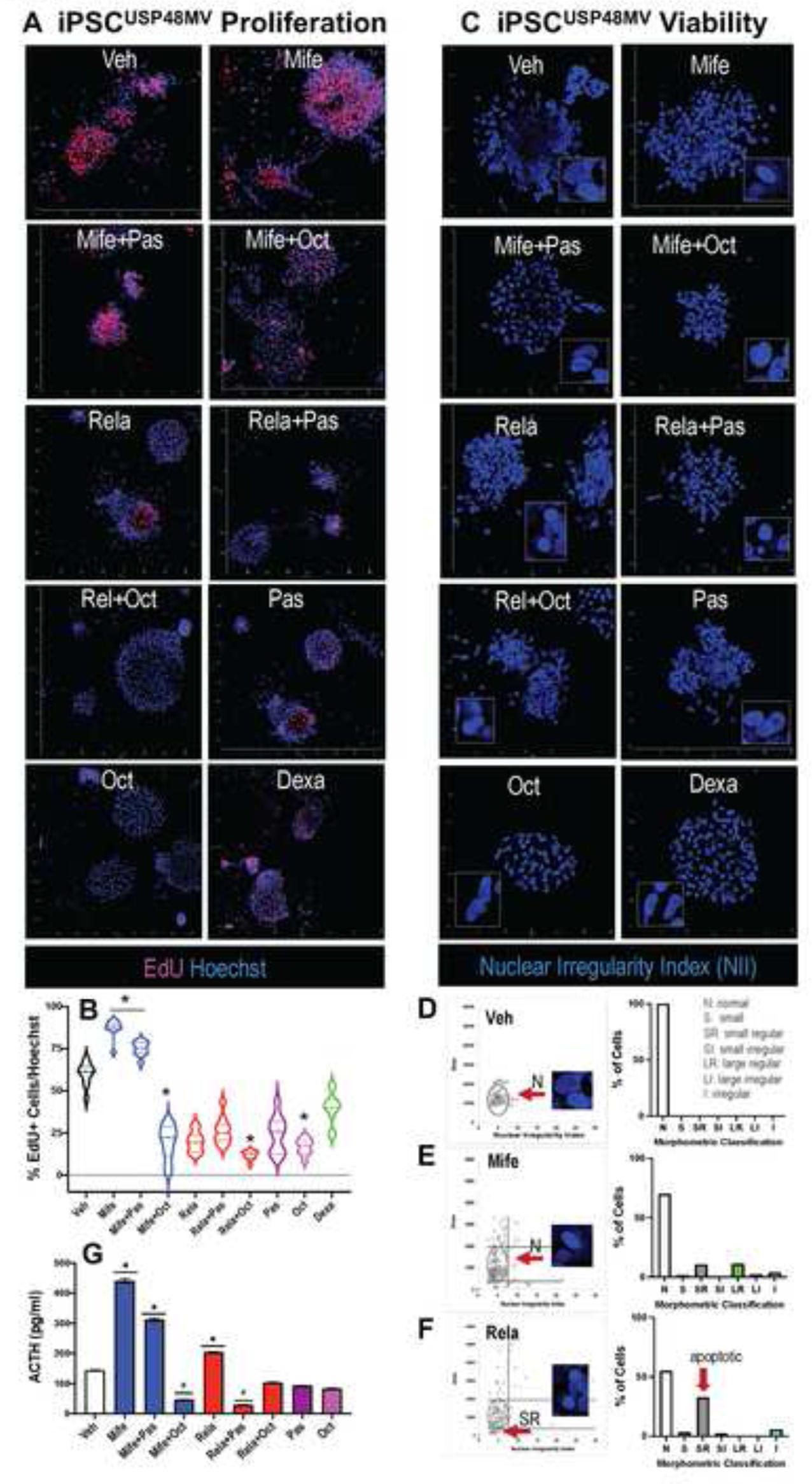
Changes in pituitary tumor cell proliferation, viability and ACTH secretion in iPSC^USP48MV^ organoids in response to mifepristone and relacorilant. **(A)** Immunofluorescence images of EdU expression in iPSC^USP48MV^ organoids in response to vehicle (Veh), mifepristone (Mife), pasireotide (Pas), octreotide (Oct), relacorilant (Rela), Mife+Pas, Mife+Oct, Rela+Pas, Rela+Oct, dexamethasone (Dexa). **(B)** Quantification of EdU positive cells of iPSC^USP48MV^ and mutant organoids. *p< 0.05 compared to iPSC^USP48MV^ organoids, n = 4 individual organoids quantified per culture. **(C)** Representative Hoescht staining of iPSC^ctrl^ organoids in response to experimental treatments for the calculation of nuclear irregularity index (NII). Nuclear morphometric analysis of iPSC^USP48MV^ organoids in response to experimental treatments with quantification shown for **(D)** Veh, **(E)** Mife, or **(F)** Rela treatments. Morphometric classification of NII was based on the normal (N), small (S), small regular (SR), small irregular (SI), large regular (LR), large irregular (LI) and irregular (I) nuclear morphology. **(G)** An ELISA was performed using conditioned media collected from iPSC^USP48MV^ cultures in response to treatments for the measurement of ACTH secretion (pg/ml). *p<0.05 compared to Veh treatment, #p<0.05 compared to Mife or Rela alone, n = 4 individual organoids quantified per culture.

Heatmap dot plots obtained after tSNE analysis of iPSC^USP48MV^ organoids showed the relative expression of the SSTR2, SSTR5, ACTH and zombie positive cells in the different phenotypic clusters in response to Rela and Mife (**Figure 7**). Quantification based on the gated identified cell clusters revealed that both Rela and Mife induced a significant increase in percentage of SSTR2 (**Figure 7A, E**) and SSTR5 (**Figure 7B, F**) positive cells. However, the magnitude of Mife induction of SSTR2 was significantly greater than that of Rela (**Figure 7A, E**). Similarly, the magnitude of Rela induction of SSTR5 was significantly greater than that of Mife (**Figure 7B, F**). In contrast to Rela, Mife led to a significant induction in the number of ACTH positive cells (**Figure 7C, G**). Rela did not significantly increase ACTH, and Pas + Rela significantly reduced ACTH expression in cultures (**Figure 7C, G**). Rela, or Rela plus Pas, clearly induced iPSC^USP48MV^ cell death as measured by the significant increase in Zombie positive cells that co-expressed ACTH and stem cell markers, a response not observed with Mife (**Figure 7D, H**).

**Figure 7:**
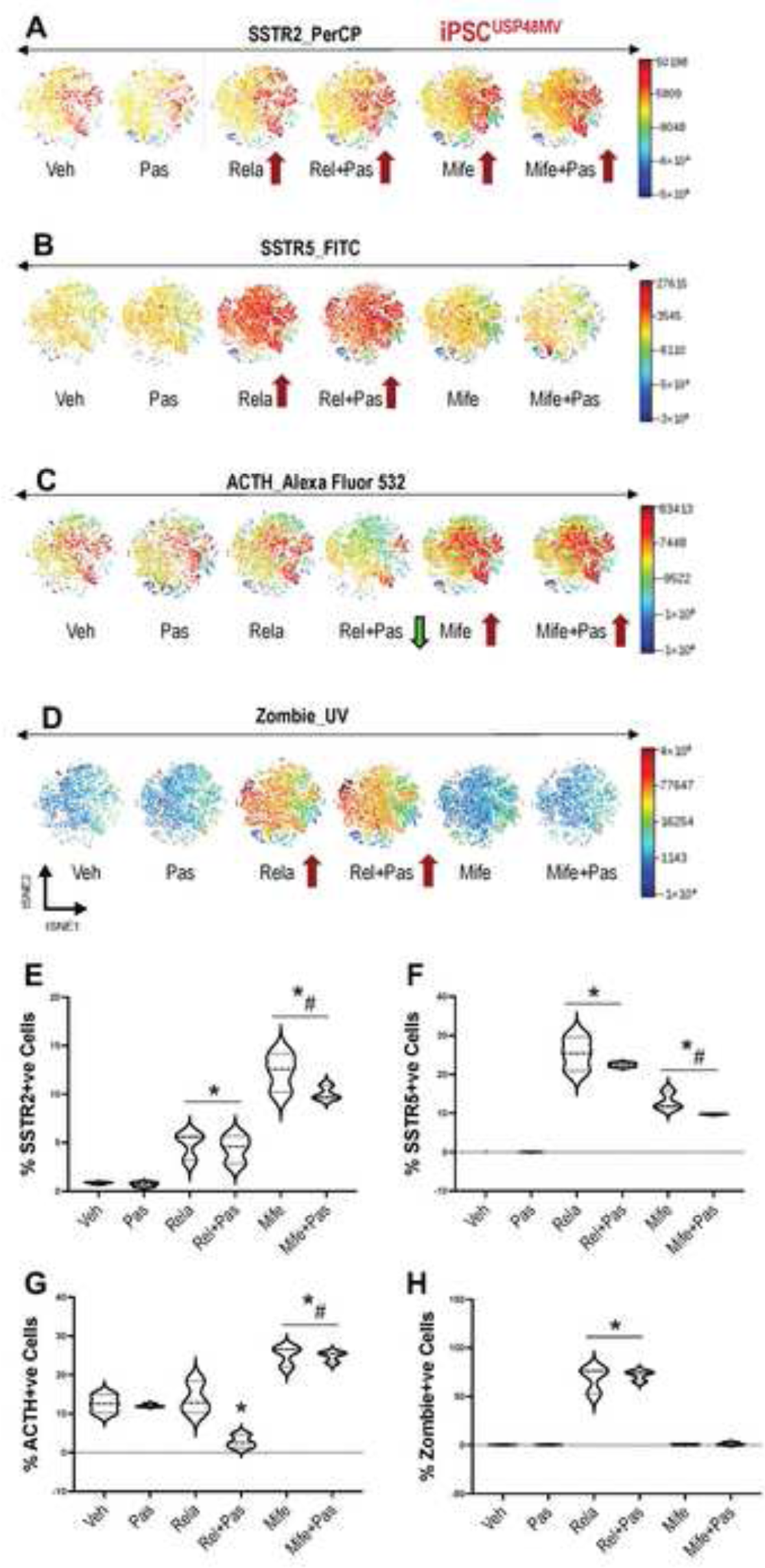
Changes in SSTR2, SSTR5 and ACTH expression, and pituitary tumor cell death in iPSC^USP48MV^ organoids in response to mifepristone and relacorilant. Fluorescent intensity of (A) SSTR2, (B) SSTR5, (C) ACTH, and (D) zombie of viSNE heatmaps for iPSC^USP48MV^ organoids in response to vehicle (Veh), pasireotide (Pas), relacorilant (Rela), Rela+Pas, mifepristone (Mife), and Mife+Pas. iPSC^USP48^ = 15000 events. Quantification of percentage of SSTR2 (E), SSTR5 (F), ACTH (G), or Zombie (H) positive cells within iPSC^USP48MV^ organoid cultures in response to experimental treatments. *p<0.05 compared to Veh.

To assess GR modulation, the GR target gene FKBP5 was measured in the PitNET organoids. While mifepristone and relacorilant significantly reduced the differential expression of FKBP5, dexamethasone caused a significant induction in gene expression (**Supplemental Figure 7A-C**). Compared to mifepristone, the magnitude of FKBP5 inhibition was significantly greater in response to relacorilant (**Supplemental Figure 7A-C**). This confirmed GR modulation in the organoid cultures in response to mifepristone and relacorilant.

Collectively, these data demonstrate that, compared to relacorilant, mifepristone preferentially induced the expression of SSTR2, ACTH secretion and PitNET organoid proliferation. In contrast, relacorilant predominantly induced SSTR5 expression and PitNET organoid apoptosis.

### 3.3. Relacorilant induces tumor cell death and reduces ACTH in combination with Pasireotide in organoids generated from CD patient PitNETs

As part of a previous study, we generated an initial biobank of organoids generated from CD patient PitNETs. Human PitNET tissue was harvested during endoscopic transsphenoidal pituitary surgery from 40 patients to generate organoids. These cultures are referred to as human PitNET derived organoids (hPITOs). **Table 1** summarizes the neuropathology reports and clinical diagnosis from cases used to generate hPITOs 37, 38, 39, 40 reported in the current study. Bright-field microscopy images of hPITOs that were generated from corticotroph PitNETs from patients diagnosed with CD revealed morphological diversity, including size, density, and regularity, among the organoid lines between individual patients and between Crooke’s cell adenoma and sparsely granulated subtypes (**Figure 8A**). Immunofluorescence confocal microscopy showed the expression of ACTH and CAM5.2 (cytokeratin 8/18) pituitary corticotroph markers (representative image of Crooke’s cell hPITO37 shown in **Figure 8B, Supplemental video 1**).

**Figure 8:**
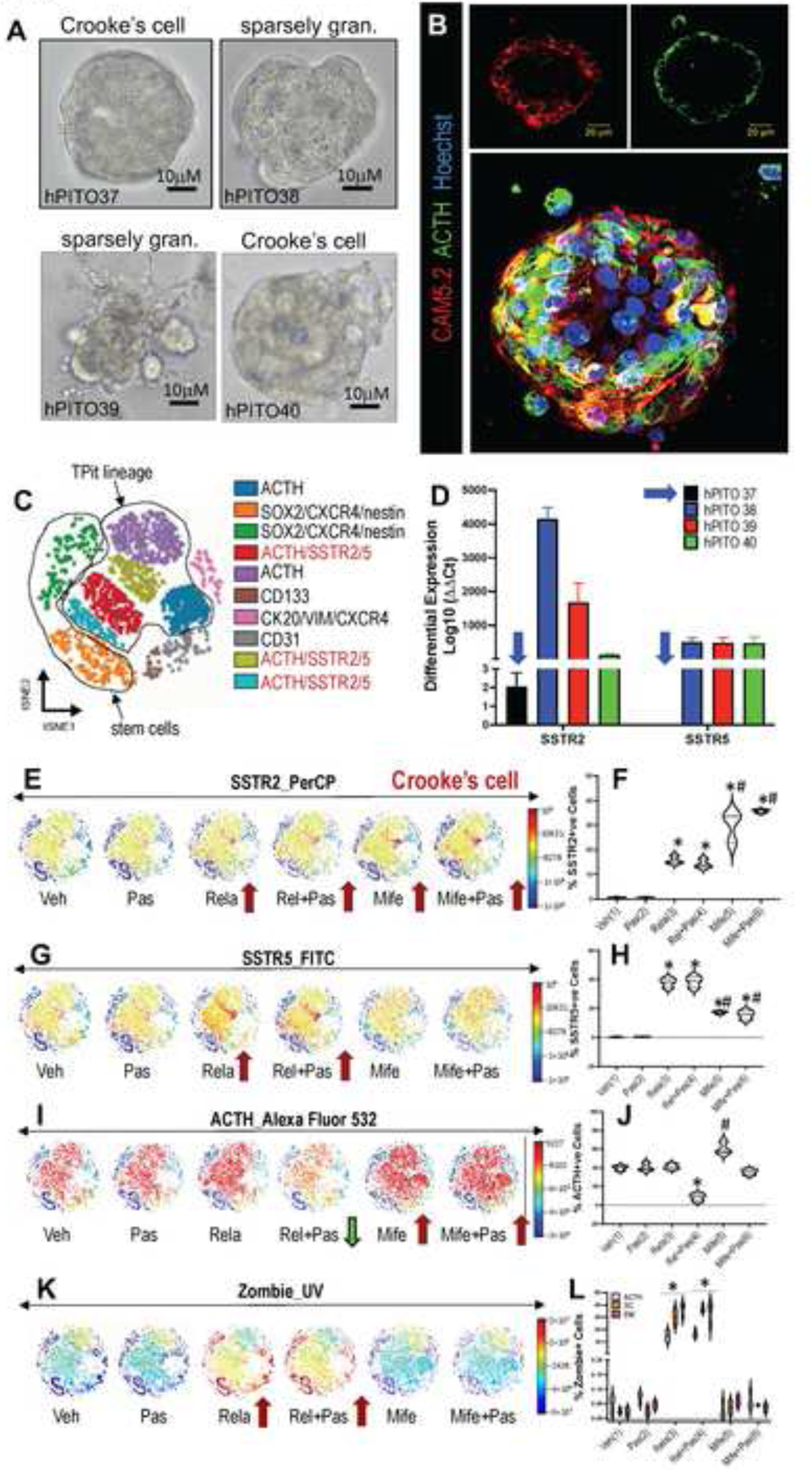
Changes in SSTR2, SSTR5 and ACTH expression, and pituitary tumor cell death in hPITOs in response to mifepristone and relacorilant. **(A)** Brightfield images of organoid cultures generated from patients with CD (hPITOs 37, 38, 39, 40). **(B)** Immunofluorescence staining using antibodies specific for CAM5.2 (red) and ACTH (green) of hPITO37. **(C)** viSNE maps showing concatenated flow cytometry standard files for hPITO37 line defining the spatially distinct cell populations using pituitary specific cell lineages, stem cell and transcription factor markers. **(D)** Differential expression of SSTR2 and SSTR5 measured in hPITO36, 37, 38, 39 and 40 lines. Fluorescent intensity of **(E, F)** SSTR2, **(G, H)** SSTR5, **(I, J)** ACTH, and **(K, L)** zombie of viSNE heatmaps for iPSC^USP48MV^ organoids in response to vehicle (Veh), pasireotide (Pas), relacorilant (Rela), Rela+Pas, mifepristone (Mife), and Mife+Pas. hPITO = 5000 events. Quantification of percentage of SSTR2, SSTR5, ACTH, or Zombie positive cells within hPITO cultures in response to experimental treatments is shown in **F, H, J and L**. *p<0.05 compared to Veh.

**Table 1:**
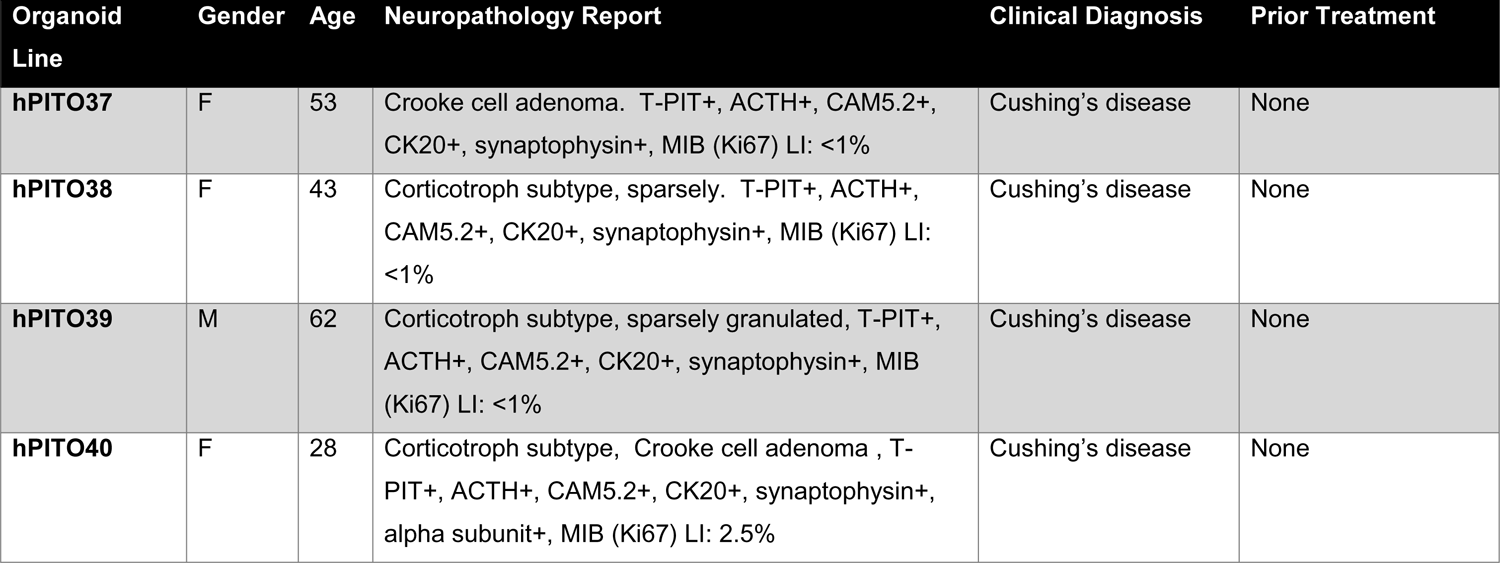
Clinical Characteristics of Pituitary Adenoma Samples Used for the Generation of Organoids

To further test the similarity in cell lineages identified between the organoid line and the patient’s tumor, we compared the immunohistochemistry from the neuropathology report to the expression pattern of pituitary tumor specific markers that were measured using Cytek™ Aurora spectral flow cytometry (**Figure 8C**). Flow cytometric analysis using Cytobank revealed that hPITOs derived from patients with CD expressed increased stem and progenitor cell markers including CXCR4, SOX2, and CD133 (**Figure 8C**). Using a gating strategy for the TPit positive cells, we identified ACTH positive cell populations co-expressing SSTR2 and SSTR5 (**Figure 8C**). Diversity in the differential expression of SSTR2 and SSTR5 among the different hPITO lines was observed when SSTR2 and SSTR5 gene expression was measured (**Figure 8D**).

Organoid line hPITO37 (Crooke’s cell adenoma subtype) expressed significantly lower SSTR2 and SSTR5 expression levels compared to the other cultures (**Figure 8D**) and was insensitive to Pas (**Supplemental Figure 3**). Therefore, we used the hPITO line to identify the differential effects of Mife and Rela on receptor expression and cell viability. Heatmap dot plots obtained after tSNE analysis of hPITO cultures showed the relative expression of the SSTR2, SSTR5, ACTH and zombie positive cells in the different phenotypic clusters in response to Rela and Mife (**Figure 8E-L**). Quantification based on the gated identified cell clusters revealed that both Rela and Mife induced a significant increase in percentage of SSTR2 and SSTR5 positive cells (**Figure 8E-H**). However, the magnitude of Mife induction of SSTR2 was significantly greater than that of Rela (**Figure 8E, F**). While both Rela and Mife induced a significant increase in percentage SSTR5 positive cells, the magnitude of Rela induction of SSTR5 was significantly greater than that of Mife (**Figure 8G, H**). In contrast, Mife significantly increased the number of ACTH expressing cells in culture, a response that was not inhibited by Pas pretreatment (**Figure 8I, J**). Rela clearly induced hPITO cell death as measured by the significant increase in Zombie positive cells in response to Rela or Rela plus Pas, a response not observed with Mife (**Figure 8K, L**). In contrast to Mife, Rela also induced significant cell death in cell populations expressing stem cell markers SOX2, CXCR4 and nestin, and an epithelial/mesenchymal hybrid cell population that co-expressed CK20, vimentin and CXCR4 (**Figure 8K, L**). Therefore, Rela sensitizes CD PitNET tissue derived organoids to pasireotide and induces cell death in several different morphologically diverse cell populations.

## 4. Discussion

Lack of effective medical therapies targeted directly to the corticotroph PitNETs is potentially attributed to the lack of human patient-relevant model systems that recapitulate the cellular complexity of the tumor. Our studies contribute to overcoming this challenge by generating a PitNET organoid model system that is applied to identifying the mechanisms of action of mifepristone and relacorilant (alone and in combination with other medical therapies for Cushing’s disease). As part of our studies, PitNET organoids were generated from: 1) CRISPR-Cas9 gene editing of patient iPSCs, and 2) CD patient corticotroph PitNETs (hPITOs). Using these advanced *in vitro* culture systems, we demonstrated that SSTR2 and SSTR5 are targets of Hedgehog transcription effector Gli1 and this response is attenuated by activation of the GR pathway. In these experiments, we have shown that mifepristone and relacorilant exerted differential effects on the induction of SSTR2 and SSTR5 expression, ACTH secretion and PitNET organoid proliferation and apoptosis.

While both mifepristone and relacorilant significantly induced the expression of SSTR2 and SSTR5, the magnitude of induction of these receptors was different. While mifepristone predominantly induced SSTR2 expression, relacorilant predominantly induced SSTR5. The SSTRs expressed on the surface of corticotroph PitNETs have often been targeted for the treatment of CD [19]. Multiple studies suggest that SSTR5 is consistently overexpressed in corticotroph PitNETs [19], however, treatments with somatostatin analogues (octreotide and pasireotide) are designed without considering variations in receptor subtype expression among individual patients with CD. We clearly show the presence of variation in the SSTR2 and SSTR5 expression amongst individual patients with CD (**Figure 8**), and, importantly, SSTR5 was significantly upregulated by relacorilant and subsequently sensitized organoid tumor cells to suppression of ACTH by pasireotide (**Figure 8**). Our findings are consistent with the knowledge that pasireotide has high-affinity binding to SSTR5 [20]. In fact, pasireotide monotherapy normalizes cortisol in up to 42% of patients with CD but assumes that all patients express this receptor subtype [21]. Combination treatment using pasireotide administered together with cabergoline and ketoconazole may increase the efficacy of the somatostatin analogue, but there is no clear consideration for changes in SSTRs during treatment [21]. Therefore, our findings suggest that treating patients with CD with relacorilant may increase the efficacy of pasireotide (or, potentially, endogenous somatostatin) suppression of ACTH secretion from corticotroph PitNETs.

In our studies we showed that Hedgehog (Hh) signaling mediates SSTR2 and 5 expression that is induced by mifepristone and relacorilant through a non-canonical Hh signaling pathway. In the iPSC organoids, GLI inhibitor GANT61 inhibited both mifepristone and relacorilant stimulated SSTR2 and SSTR5. The SMO inhibitor ketoconazole (which also inhibits cortisol synthesis) failed to block the induction of both SSTR2 and SSTR5 by both compounds. Upon ligand binding to the transmembrane receptor Patched-1 (PTCH1), the repression of PTCH1 on transmembrane transducer SMO is released. De-repressed SMO triggers the activation of GLI zinc finger transcription factors GLI1, GLI2 and GLI3. Activated GLI1 protein translocates into the nucleus, where several target genes are regulated [42]. Our studies suggest that within the corticotroph PitNETs, SSTR2 and SSTR5 are transcriptional targets of GLI. The failure of the SMO inhibitor ketoconazole to inhibit mifepristone and relacorilant induction of SSTR expression suggests that the signaling pathway is non-canonical (**Supplemental Figure 8**). Non-canonical Hh signaling pathways have been classified as 1) Type I pathways involving signaling through PTCH1 independent of SMO, 2) Type II pathways involving signaling through SMO independent of GLI, and 3) SMO-independent activation of GLI such as that observed in the current study [42, 43]. In support of our findings, crosstalk between GLI and GR (NR3C1) signaling has been reported in T cell acute lymphoblastic leukemia [44]. In addition, studies in the adult stomach demonstrated that the Hh pathway is critical for the regulation of somatostatin and SSTR signaling [45, 46]. While Hh signaling is known to be essential during the embryonic development of the pituitary and in the adult gland [47], this is the first report demonstrating that Hh signaling via Gli1 regulates the expression of SSTR2 and SSTR5 in PitNET organoids.

In contrast to mifepristone, relacorilant induced PitNET cell death in treated organoid cultures without significant stimulation of ACTH secretion. In concurrence with the organoid *in vitro* studies, tumor regression in two patients with macroadenomas treated with relacorilant for three months have been reported in an initial study based on standard of care imaging of the pituitary gland [17]. This unexpected finding is further being investigated in an ongoing Phase 3 study of relacorilant in patients with Cushing’s syndrome (GRACE Study, <u>NCT03697109</u>). Such macroadenoma regressions have not been observed with mifepristone. Based on the negative feedback mechanism regulating the hypothalamic-pituitary-adrenal (HPA) axis, we propose the following mechanisms to explain the differential effects between mifepristone and relacorilant. First, mifepristone predominantly induced SSTR2 expression. However, unlike relacorilant, mifepristone treatment significantly increased ACTH secretion from PitNET both *in vivo* and *in vitro* (**Supplemental Figure 8)**. Our organoid experiments are supported by patient data showing that mifepristone increases ACTH [13, 14], with a concomitant increase in cortisol secretion without tumor growth [19, 48]. In contrast, relacorilant predominantly induced SSTR5 expression without an increase in ACTH secretion and tumor cell death (**Supplemental Figure 8**). A plausible explanation for PitNET regression in response to relacorilant treatment may be the known heterodimerization between SSTR2 and SSTR5 that occurs following the selective activation of SSTR2 but not human SSTR5 or concurrent stimulation [49, 50]. Heterodimerization between SSTR2 and SSTR5 leads to an increased recycling rate and a greater propensity of SSTR2 to induce tumor growth inhibition [50, 51]. Thus, *in vivo*, the significant effect of mifepristone on SSTR2 and increased receptor signaling offsets the proliferative effects of this antagonist and may explain the lack of tumor growth in treated patients. Relacorilant may also induce PitNET apoptosis independently of SSTR activation (**Supplemental Figure 8**). In support of this notion, studies in several cancers that are driven by the dysregulation of these receptors, including leukemia, breast, prostate, lung and ovarian cancers, show promise of targeting the GR as a strategy for combination treatment [52, 53]. In addition, relacorilant induces apoptosis *in vitro* using pancreatic and ovarian cancer cell lines [54]. The pro-apoptotic effects of GR modulators are tissue specific and may be a result of targeting the apoptotic/cycle signaling pathways and genes such as *Bcl2* [55]. The mechanism of action of relacorilant on PitNET cell apoptosis requires further investigation. Collectively, these studies support broader investigation of relacorilant alone or in combination with somatostatin analogs as pre- or post-operative medical treatment in patients with CD. While many investigators have proposed using organoids in personalized medicine for the targeted treatment of cancers [29, 56–58], this study is the first to execute this approach for the potential treatment of corticotroph type PitNETs associated with the development of CD. Existing medical therapy for CD remains suboptimal with negative impact on health and quality of life, including the considerable risk of therapy resistance and tumor recurrence. Our data demonstrate that organoids derived from corticotroph type PitNETs consist of differentiated cell lineages, stem/progenitor cells, and stromal cells that replicate much of the patient’s own tumor pathology and function clearly documented by extensive high-throughput flow cytometry. Previously published *in vitro* experiments in the field of CD research were performed using pituitary cell lines or spheroids, aggregates and tumoroids that do not replicate the primary pituitary tumor microenvironment due to cell transformation and/or unphysiological 2D culture conditions [59–61]. Pituitary research has largely been conducted using cell culture techniques using rat (GH3) or mouse (AtT20) pituitary-like cell lines lacking a multicellular identify reflective of the PitNET [62, 63]. Recent advances have led to the development of pituitary tissue generated organoids, but these are limited to the use of transgenic mouse models as the source [64–66]. Nys *et al.* [67] reported the generation of human pituitary tumor organoids from a single stem cell as claim of true organoids due to the clonality. Unlike our studies, the multicellular complexity was not validated by the protein expression or hormone secretion from pituitary cell lineages in these cultures [67]. It is fundamental to note that according to the National Cancer Institute (NCI, NIH), an ‘organoid’ is defined as “A tiny, 3-dimensional mass of tissue that is made by growing stem cells (cells from which other types of cells develop) in the laboratory.” [68]. Our cultures begin with both single and 3-4 cell clusters isolated from the native CD patient PitNET tissue that harbor the stem cells and begin a process of ‘budding’, growth and differentiation as documented by **supplemental videos 2 and 3** and the comprehensive spectral flow cytometry analysis documenting functional cell lineages. Our PitNET organoids are consistent with gastrointestinal tissue derived organoids including that begin from cell clusters, crypts or glands [29, 69, 70]. Importantly, the videos show a different process than what would be an expected observation for the formation of a tumoroid which is the migration and adhesion of initially separate cells to form an aggregate. There are the reports of mouse nonadherent spheres with stem/progenitor characteristics [71], and human embryonic stem cell generated spheroids or patient derived tumoroids that also lack a multicellular identify and consist of poorly differentiated cells [72–77]. Therefore, the findings reported here are significant because we offer an advanced *in vitro* approach that reveals cell populations expressing stem cell markers that potentially contribute to the support of tumor growth and targeted to prevent tumor recurrence. The ability of relacorilant to induce PitNET cell death in a multicellular culture system is the first step in the development of effective targeted therapies for patients with CD and potentially, other PitNET subtypes.

## SUPPLEMENTAL VIDEOS

**Supplemental Video 1: 3D rendering of a confocal image captured from CD PitNET tissue derived organoid culture.** Movie of z-stack captured through the hPITO37 immunofluorescently stained for CAM5.2 (red) and ACTH (green). Hoechst was used for nuclear staining (blue).

**Supplemental Video 2: Time-lapse video of iPSC PitNET derived organoid culture (area 1).** Video of live time-lapse brightfield images, captured by a Nikon ECLIPSE Ti2 microscope, of organoid culture scanned automatically every 30 minutes for 72 hours.

**Supplemental Video 3: Time-lapse video of iPSC PitNET derived organoid culture (area 2).** Video of live time-lapse brightfield images, captured by a Nikon ECLIPSE Ti2 microscope, of organoid culture scanned automatically every 30 minutes for 72 hours.

## SUPPLEMENTAL FIGURE LEGENDS

**Supplemental Figure 1: Generation of pituitary tumor organoids from iPSCs or adenoma tissue. (A)** Differentiation schedule for the generation of pituitary organoids derived from iPSCs expressing somatic mutations driving the development of CD. Pituitary organoids were also generated based on the outlined schedule using iPSCs generated from PBMCs collected from a patient with CD, or a healthy individual. Bright field images demonstrating morphological variation is observed between iPSC^ctrl^ and iPSC^USP48^ or iPSC^USP8^ lines and organoids. **(B)** Overview of the generation of hPITOs from adenoma tissues collected from patients with CD.

**Supplemental Figure 2: CRISPR mutation design and validation of gene edited iPSC generated organoids. (A)** Design of somatic mutations associated with CD (USP8, USP48, and BRAF) in corticotroph pituitary tumors. Restriction Fragment Length Polymorphism (RFLP) validation was used to confirm successful gene editing. Undigested (U) DNA **(B)** were compared to samples digested with the restriction enzyme (D). Respective restriction enzymes for each mutation were used (AvaII for BRAF, SpeI for USP48, BstBI for USP8). **(C)** Arrows indicate the U and D bands on the gel. iPSC^Ctrl^ displayed no separation of bands in digested versus undigested samples or amplification of the mutated genes.

**Supplemental Figure 3: Drug responses of hPITOs and adjacent normal pituitary organoids.** Dose-response curves using hPITO36-40 and hPITO37N and 38N generated from adjacent normal pituitary tissue in response **(A, B)** pasireotide, **(C, D)** mifepristone, and **(E, F)** relacorilant.

**Supplemental Figure 4: Expression pattern of major hormone-producing cell lineages in iPSCs differentiated to pituitary organoids.** Expression of PIT-1 (green), ACTH (green), GH (red), FSH (red), LH (green), PRL (red) and synaptophysin (synapto, green) with co-stain Hoechst (nuclei, blue) was measured by immunofluorescence using chamber slides collected at day 4 (D4) of the differentiation schedule of **(A)** control iPSCs (iPSC^ctrl^) and iPSCs expressing **(B)** USP48MV (iPSC^USP48MV^), **(C)** USP48MI (iPSC^USP48MI^) and **(D)** USP8 (iPSC^USP8^) mutations. Red arrows highlight the increased expression of ACTH and synaptophysin with the concomitant loss of PIT1, GH, FSH, LH and PRL in iPCSs expressing somatic mutations USP48 and USP8. Quantification of the percentage of positive cells is shown in the dot plots for each iPSC line.

**Supplemental Figure 5: Nuclear Morphometric Analysis of iPSC^ctrl^ organoid cultures in response to mifepristone and relacorilant.** Morphometric classification of NII was based on the normal (N), small (S), small regular (SR), short irregular (SI), large regular (LR), large irregular (LI) and irregular (I) nuclear morphology. Distribution of nuclear morphology in response to in response to vehicle (Veh), mifepristone (Mife), pasireotide (Pas), octreotide (Oct), relacorilant (Rela), Mife+Pas, Mife+Oct, Rela+Pas, Rela+Oct, dexamethasone (Dexa) was plotted.

**Supplemental Figure 6: Nuclear Morphometric Analysis of iPSC^USP48MV^ organoid cultures in response to mifepristone and relacorilant.** Morphometric classification of NII was based on the normal (N), small (S), small regular (SR), short irregular (SI), large regular (LR), large irregular (LI) and irregular (I) nuclear morphology. Distribution of nuclear morphology in response to in response to vehicle (Veh), mifepristone (Mife), pasireotide (Pas), octreotide (Oct), relacorilant (Rela), Mife+Pas, Mife+Oct, Rela+Pas, Rela+Oct, dexamethasone (Dexa) was plotted.

**Supplemental Figure 7: Changes in FKBP5 expression in response to mifepristone, relacorilant and dexamethasone.** Quantitative RT-PCR was performed on RNA extracted from **(A)** iPSC^ctrl^, **(B)** iPSC^USP48MV^, and **(C)** iPSC^USP8^, organoids treated with vehicle (Veh), mifepristone (Mife), relacorilant (Rela) or dexamethasone (Dexa).

**Supplemental Figure 8: Summary of divergent effects of mifepristone and relacorilant on SSTR2 and SSTR5 expression, ACTH secretion, and pituitary tumor proliferation and apoptosis.** Mifepristone induces SSTR2 with greater magnitude compared to SSTR5. Relacorilant predominantly induces SSTR5 and tumor cell apoptosis. GLI signaling inhibitor GANT61 blocks SSTR2 and SSTR5 expression in response to mifepristone and relacorilant. SMO inhibitor ketoconozaole has no effect on the induction of SSTR2 and SSTR5 by GR modulators mifepristone and relacorilant. GR antagonists are likely to signal via a non-canonical GLI1 SMO-independent signaling pathway. Arrows indicate the magnitude of induction.

## ACKNOWLEDGEMENTS

This research was supported by Department of Cellular and Molecular Medicine (University of Arizona College of Medicine) startup funds (Zavros). We acknowledge Maga Sanchez in the Tissue Acquisition and Cellular/Molecular Analysis Shared Resource (TACMASR University of Arizona Cancer Center) for assistance with embedding and sectioning of organoids. We would also like to acknowledge Patty Jansma (Marley Imaging Core, University Arizona) and, Douglas W Cromey (TACMASR imaging, University of Arizona Cancer Center) for assistance in microscopy. Research reported was partly supported by the National Cancer Institute of the National Institutes of Health under award number P30 CA023074 (Sweasy). We would like to acknowledge the assistance of Tina Schlafly (Corcept) for help with manuscript editing. Finally, the authors thank the patients who consented to donate tissues and blood for the development of the organoids. Without their willingness to participate in the study, this work would not be possible.

## AUTHOR CONTRIBUTIONS

**Saptarshi Mallick:** Conceptualization, Methodology, Software, Data curation, Writing-Original draft preparation, Software, Formal Analysis, Reviewing and Editing; **Jayati Chakrabarti**: Methodology, Software, Data curation, Writing – Original draft preparation, Validation, Formal Analysis, Reviewing and Editing; **Jennifer Eschbacher:** Resources, Validation, Reviewing and Editing; **Andreas G. Moraitis:** Conceptualization, Resources, Reviewing and Editing; **Andrew Greenstein:** Conceptualization, Resources, Reviewing and Editing; **Jared Churko:** Methodology, Validation, Formal Analysis, Reviewing and Editing; **Kelvin W. Pond:** Methodology, Validation, Formal Analysis, Reviewing and Editing**; Antonia Livolsi:** Methodology; **Curtis Thorne:** Resources; **Andrew S. Little:** Conceptualization, Writing – Original draft preparation, Validation, Reviewing and Editing, Resources, Visualization; **Kevin Yuen:** Conceptualization, Writing, Original draft preparation, Validation, Reviewing and Editing, Resources, Visualization; **Yana Zavros:** Conceptualization, Methodology, Software, Data curation, Writing – Original draft preparation, Software, Validation, Formal Analysis, Reviewing and Editing, Resources, Visualization, Investigation, Supervision.

## Abbreviations

ACTH: Adrenocorticotropin Hormone

BMP4: Bone Morphogenetic Protein 4

CD: Cushing Disease

CDH23: Cadherin Related 23

DMEM/F12: Dulbecco’s Modified Eagle Medium/Nutrient Mixture F-12

DMSO: Dimethylsulfoxide

DPBS: Dulbeccos Phosphate Buffered Saline

ELISA: Enzyme-linked Immunoassay

FSH: Follicle-Stimulating Hormone

GH: Growth Hormone

GLI1: GLI family zinc finger1

GR: Glucocorticoid Receptor

hPITO: Human Pituitary Organoids

iPSC: Induced Pluripotent Stem Cells

MEN1: Multiple endocrine neoplasia link type 1

Mife: Mifepristone

NII: Nuclear Irregularity Index

NMA: Nuclear Morphometric Analysis

Oct: Octreotide

Pas: Pasireotide

PBMCs: Peripheral Blood Mononuclear Cells

PBST: PBS with Tween™ 20

PCR: Polymerase Chain Reaction

PIT1: Pituitary-specific positive transcription factor 1

PitNET: Pituitary Neuroendocrine Tumor

PRL: Prolactin

PTCH1: Protein patched homolog 1

Rela: Relacorilant

SFDM: Serum-free Defined Media

SHH: Sonic Hedgehog

SMO: Smoothened

SSTR: Somatostatin Receptor

TPit: T-Box Transcription Factor 19

USP48: Ubiquitin carboxyl-terminal hydrolase 48

USP8: Ubiquitin carboxyl-terminal hydrolase 8

